# Can Intestinal Absorption of Dietary Protein Be Improved Through Early Exposure to Plant-Based Diet?

**DOI:** 10.1101/2020.01.24.917856

**Authors:** Giovanni S. Molinari, Vance J. McCracken, Michal Wojno, Simona Rimoldi, Genciana Terova, Karolina Kwasek

## Abstract

Nutritional Programming (NP) has been studied as a means of mitigating the negative effects of dietary plant protein (PP), but the optimal timing and mechanism behind NP are still unknown. The objectives of this study were: 1) To determine whether zebrafish (*Danio rerio*) can be programmed to soybean meal (SBM) through early feeding and broodstock exposure to improve SBM utilization; 2) To determine if NP in zebrafish affects expression of genes associated with intestinal nutrient uptake; 3) To determine if early stage NP and/or broodstock affects gene expression associated with intestinal inflammation or any morphological changes in the intestinal tract that might improve dietary SBM utilization. Two broodstocks were used to form the six experimental groups. One broodstock group received fishmeal (FM) diet (FMBS), while the other was fed (“programmed with”) SBM diet (PPBS). The first (**(+) Control**) and the second group (**(-) Control**) received FM and SBM diet for the entire study, respectively, and were progeny of FMBS. The last four groups consisted of a non-programmed **(FMBS-X-PP** and **PPBS-X-PP)** and a programmed group **(FMBS-NP-PP** and **PPBS-NP-PP)** from each of the broodstocks. The programming occurred through feeding with SBM diet during 13-23 dph. The non-control groups underwent a PP-Challenge, receiving SBM diet during 36-60 dph. During the PP-Challenge, both PPBS groups experienced significantly lower weight gains than the (+) Control group. NP in early life stages significantly increased the expression of PepT1 in PPBS-NP-PP, compared to PPBS-X-PP. NP also tended to increase the expression of *fabp2* in the programmed vs. non-programmed groups of both broodstocks. The highest distal villus length-to-width ratio was observed in the dual-programmed group, suggesting an increase in surface area for nutrient absorption within the intestine. The results of this study suggest that NP during early life stages may increase intestinal absorption of nutrients from PP-based feeds.

## Introduction

Plant protein- (PP) based diets in aquaculture are advancing but, are not yet fully suitable replacements for fishmeal (FM) for many species due to their adverse effects on growth and health. Soybean meal (SBM) is the most commonly used protein source in plant-based diets but, has been shown to reduce fish growth at high dietary inclusion levels (above 25%) and to cause inflammation in the intestines of many fish species due to presence of anti-nutritional factors [1]. The reduced growth performance of fish fed SBM-based diets has been studied in both carnivorous and omnivorous species [2–4]. For example, rainbow trout (*Oncorhynchus mykiss)* fed a PP-based diet showed a 17% decrease in growth and a significant reduction in feeding efficiency [2]. Wang et al. [4] concluded that, in orange-spotted grouper (*Epinephelus coioides*), increasing levels of dietary soybean meal led to reduced growth performance and increased intestinal damage. Similar results were observed in the omnivorous species, Nile tilapia (*Oreochromic niloticus*). Diets tested included 0, 33, 66, and 100% inclusion of PP; increasing levels of PP inclusion resulted in corresponding decreases in growth rate [3]. Reduced growth performance of fish fed PP-based diets is a key restriction on the use of PP in aquaculture. Many lower quality plant protein-based diets are not yet able to match the growth performance and feeding efficiency of fishmeal or high-quality protein sources (i.e. wheat/corn gluten, soy protein concentrate), restricting their ability to fully replace them in aquaculture.

Nutrient absorption can be studied by measuring the expression of certain genes within the intestine. PepT1, also known as *slc15a1* [5], is an intestinal protein transporter responsible for peptide uptake [6]. Increased expression of intestinal peptide transporters is a key factor in the absorption of feed components in fish [6]. Plant protein sources have a negative effect on PepT1 expression [7, 8]. A dietary inclusion of just 15% green pea meal significantly reduced PepT1 expression in European sea bass (*Dicentrarchus labrax*), correlating with a reduced growth performance compared to FM-fed fish [7]. Similar results were observed on sea bream (*Sparus aurata*), where a 15% FM replacement with PP led to reduced PepT1 expression and decreased growth [8]. Both studies provide support that intestinal PepT1 expression is a good marker for absorption efficiency and protein quality within feeds [7, 8]. Furthermore, SBM causes a down-regulation of *fabp2* – a fatty-acid binding protein responsible for lipid absorption within the gut [9] causing a disruption of the epithelial transport of lipids and their reduced absorption [9, 10]. Decreased expression of *fabp2* has been observed particularly in fish that experienced SBM-induced inflammation in the distal portion of the intestine [5].

Intestinal inflammation has been assessed in previous studies, using qPCR to measure the regulation of inflammatory markers within the intestines. In a study done on larval zebrafish (*Danio rerio*), the pro-inflammatory cytokine tumor necrosis factor-α (*tnfa*) was measured in order to analyze the intestinal health of zebrafish fed soy-based feeds [5]. Expression of *tnfa* was significantly higher in fish fed SBM than in fish fed a standard fishmeal diet, representing increased intestinal inflammation [5]. In Atlantic salmon (*Salmo salar*), intestinal expression of *il17a*, *il1b*, and other genes that are also considered pro-inflammatory markers were increased in fish exposed to SBM [11]. In that study, the increased expression of those inflammatory markers was observed within three days of exposure, showing that the process of intestinal inflammation occurs rather quickly in response to dietary SBM [11]. Furthermore, exposure to SBM is associated with shortened intestinal mucosal folds, thickened lamina propria, and an increased number of goblet cells [10, 12]. Those histological responses are consistent with intestinal inflammation [12] and are responsible for reduced feed utilization efficiency and poor growth performance [1].

One approach to improving the utilization of plant-based feeds in aquaculture is Nutritional Programming (NP). NP is the concept that an organism can be ‘programmed’ to a certain diet through exposure to that diet during the early stages of development. More specifically, if fish are exposed to the PP during development, they are then able to adapt to that protein source and process plant-based feed better during later life stages. Previous studies have shown that early feeding of plant-based diets can increase feed acceptance and utilization in fish fed the same plant-based feed later in life [13]. For example, rainbow trout programmed with early (juvenile) feeding of dietary PP had a 42% higher growth rate and 30% higher feed intake during PP feeding later in life compared to fish that had not received plant-based diets during development [13]. In addition to the improved growth response related to NP, there is evidence to support the use of NP to help mitigate intestinal inflammation associated with SBM feeds. Perera and Yufera [5], found that certain features of intestinal inflammation could be programmed through the early feeding of SBM-based feeds. However, more knowledge regarding the relationship between NP and intestinal responses in fish is still needed to better understand the mechanism of NP and its role in improving dietary PP utilization.

NP mode of action occurs through the epigenome. The epigenome of an individual is what programs its genome and affects the phenotypic expression of a given genomic sequence [14]. Environmental stimuli can trigger epigenetic marks that affect the expression of certain genes [15]. The epigenetic effect can last multiple generations, meaning it has the possibility to impact the phenotype of the broodstock and also the phenotypes of their offspring [15]. Diet and nutrients within the diet are environmental stimuli that can change the epigenome so that specific genes, for example within the digestive tract, could be up- or down-regulated as a response to the diet; these changes in turn can be inherited by the offspring [15]. Broodstock programming is feasible in many fish species in which large part of larval development occurs during the yolk-sac stage, where the yolk, the composition of which is greatly influenced by parental diet, is the only source of nutrition during endogenous feeding [16, 17]

If digestive processes in broodstock are able to adapt to the plant-based diets through altered expression of specific genes associated with nutrient transportation or inflammatory response, then those adaptations could potentially be passed down to offspring through the epigenome. Studies testing the effects of feeding gilthead seabream (*Sparus aurata*) broodstock with low, moderate, and high fish oil (FO) and FM diets found that feeding moderate FO/FM levels improved growth and feeding efficiency of their respective offspring that were fed a low FO/FM diet [18]. That study utilized rapeseed meal and linseed oil as replacements for FO and FM, so it is important to determine if similar effects can be observed using SBM in different species.

Due to a relatively small amount of research, the mechanism behind NP is still unknown. We hypothesize, however, that improved growth performance of fish fed SBM-based diet during adulthood results from adaptation of the intestinal tract to that particular protein source when exposed to it during broodstock or early developmental stages. The objectives of this study were: 1) To determine whether fish can be programmed to dietary SBM through broodstock feeding and in combination with NP of the early stages to improve SBM utilization in the offspring during the grow-out phase; 2) To determine if NP affects expression of genes associated with intestinal inflammation and peptide uptake, and 3) To determine if NP through broodstock feeding and in combination with NP of early stages induces any morphological changes in the intestinal tract. Zebrafish, a popular model organism due to their high fecundity and fully annotated genome [19], were used for this experiment. Zebrafish can feed on vegetal and animal protein sources, so it is an ideal species to model the nutritional pathways of both carnivorous and omnivorous aquaculture species [20]. Moreover, zebrafish are a good model to study diet-induced inflammation in the intestines of aquaculture species [19].

## Materials and Methods

This study was conducted at the McLafferty Annex of Center of Fisheries, Aquaculture, and Aquatic Sciences at Southern Illinois University, Carbondale (SIUC), IL. All experiments were approved by the Institutional Animal Care and Use Committee (IACUC) of SIUC (protocol# 18-007).

All experiments were carried using recirculated aquaculture system (Pentair Aquatic Eco-systems, Cary, NC). The system consisted of 3 L and 5 L tanks, stacked in rows. Water in the system was constantly filtered and recirculated using a mat filter, UV light, a carbon filter, and biofiltration. The average water temperature was 27.1 ± 0.2°C, the pH was 7.28 ± 0.72, and the salinity was kept between 1 and 3 ppt (higher during the first feeding to prolong the viability of the live food [21]). The photoperiod consisted of 14 hours of darkness and 10 hours of light, with the overhead lights on 8:00-18:00.

### Feed Preparation and Formulation

Dry components of feeds were ground to a fine particle size (0.5-0.25 mm) using a centrifugal mill (Retsch 2M 100, Haan, Germany). Once ground, components were mixed (Farberware Mixer, Fairfield, CA) to achieve uniform dispersion of all ingredients within the mix. After mixing, feeds were forced through an extruder (Caleva Extruder 20, Sturminster Newton Dorset, England) and then spheronized (Caleva Multibowl Spheronizer, Sturminster Newton Dorset, England) to obtain solid, spherical pellets. All feeds were then freeze-dried (Labconco FreeZone 6, Kansas City, MO) to remove moisture. Pellets were separated by size using a vibratory sieve shaker (Retsch AS 200 Basic, Haan, Germany) to appropriate sizes. Both diets were isoenergetic and isonitrogenous. Crude protein level of the diets was 48.9%, with a crude lipid level of 10%. The SBM diet replaced 100% of fishmeal with a combination of soybean meal and soy protein isolate. Soy protein concentrate was included as necessary to adjust dietary crude protein while leaving room for other ingredients in the formulation, including a minimum level of starch to allow expansion of the experimental diets. Feed formulations for this study are listed in Table 1. In this study, the SBM diet serves as the PP-based diet.

**Table 1.**
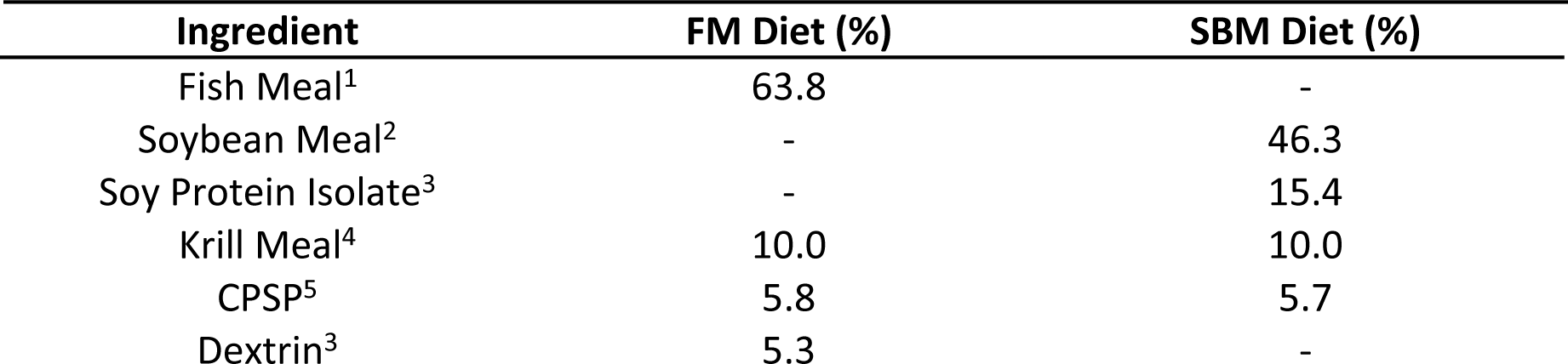
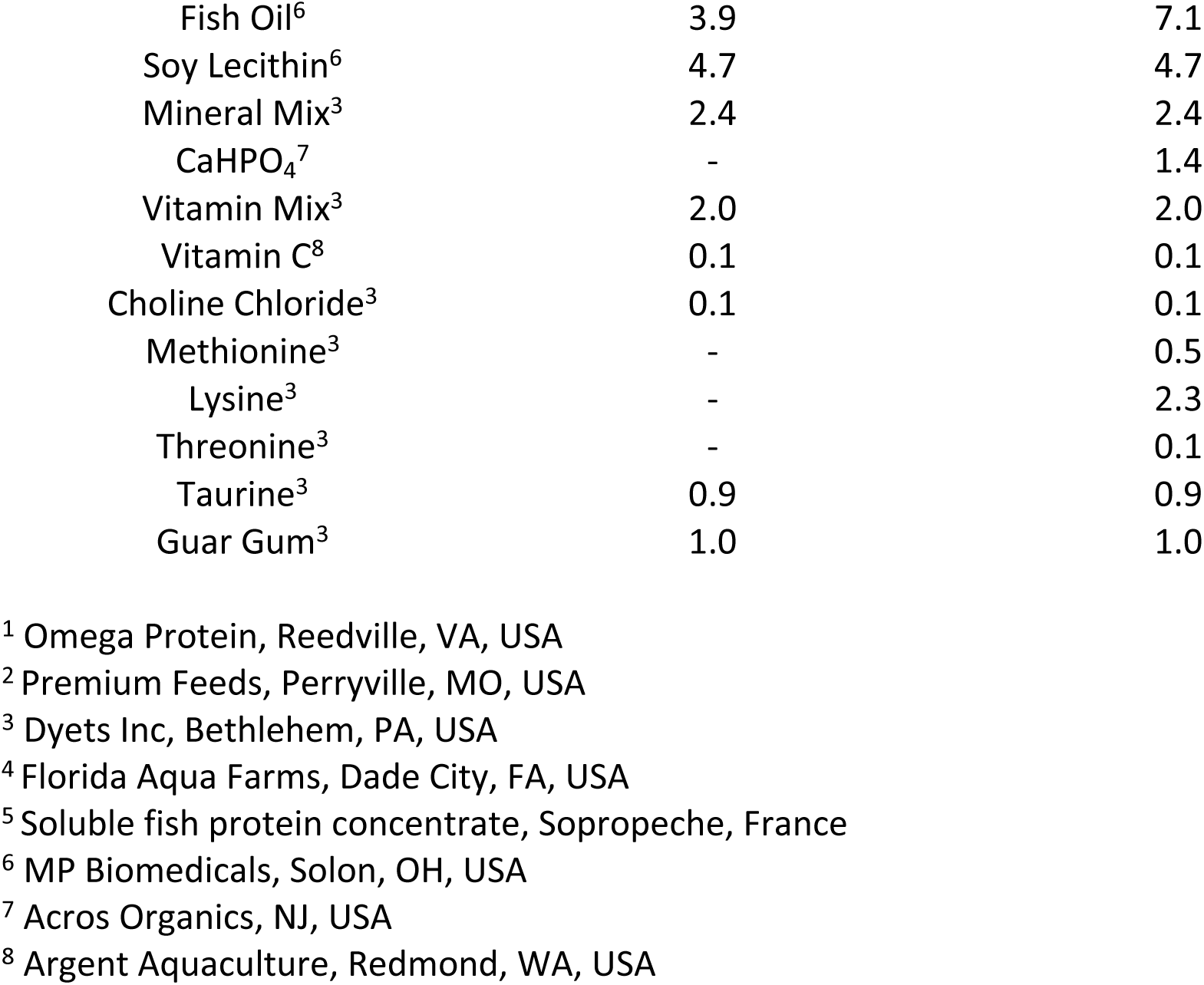
Feed formulation (g/100g) of experimental diets.

### Broodstock Rearing and Spawning

Broodstock used in this experiment were separated into a fishmeal group (FMBS) and a PP group (PPBS). Initially, males and females from each group were separated and during this period, FMBS and PPBS groups were fed twice a day with a fishmeal- and PP-based diet, respectively. Both groups also received *Artemia* nauplii (Brine Shrimp Direct, Ogden, UT) as supplemented feed twice a day prior to breeding to ensure proper gonadal development, particularly in case of PP diet-fed group. After 4 weeks, males and females in each group were combined into their respective breeding tanks. The breeding tanks included a mesh sheet at the bottom with an artificial plant to induce spawning. After 24 hours, broodstock were removed to allow offspring to hatch.

### Experimental Design and Feeding Regimen

There were total of six experimental groups in this study. The four non-control groups underwent plant-protein challenge (PP-Challenge). The PP-Challenge consisted of exposure to dietary SBM in later life stages of experimental fish (36-60 days-post hatch (dph)) to determine how each group responds to PP in terms of growth and the status of intestinal epithelial lining. Each experimental group were housed in three replicate tanks, with 70 fish in a 3 L tank. At 3 dph larvae from each broodstock group (FMBS and PPBS) were randomly distributed to their respective experimental tanks. Offspring from each broodstock group were separated into different experimental groups. Four groups came from the fishmeal-fed broodstock (FMBS): 1) **(+) Control** group – the purpose of this group was to determine the intestinal health and growth rates of fish that were continuously fed a fishmeal-based diet immediately following the initial live-feed period; 2) **(–) Control -**the purpose of this group was to observe the growth and intestinal health of fish that are only fed PP-based throughout their life immediately following the initial live-feed period; 3) **FMBS-X-PP** – this group represented how fish that were not programmed through broodstock or early exposure to PP will respond to PP Challenge; 4) **FMBS-NP-PP** – this group meant to show us how fish that are only programmed to PP in early development, but not in broodstock stage, will respond to the PP-Challenge. Due to low fecundity and hatching rate, only two experimental groups came from the PP-fed broodstock (PPBS): 5) **PPBS-X-PP**- used to determine how only programming through broodstock (and not induced in early development) will affect the fish response to the PP-Challenge; 6) **PPBS-NP-PP** – a group that underwent both broodstock and early feeding programming and represents how the combination of the two will affect growth and intestinal health during the PP-Challenge.

Experimental groups and feeding regimen are displayed in Fig 1. Fish were fed *ad libitum* from 3-26 dph and then provided restricted feeding rates from 26-60 dph. From 26-45 dph the restricted feeding rate was 9% of the total biomass per day, and from 45-60 dph the feeding rate was decreased to 8%. All feeding rates were adjusted according to fish growth and feed intake.

**Fig 1.**
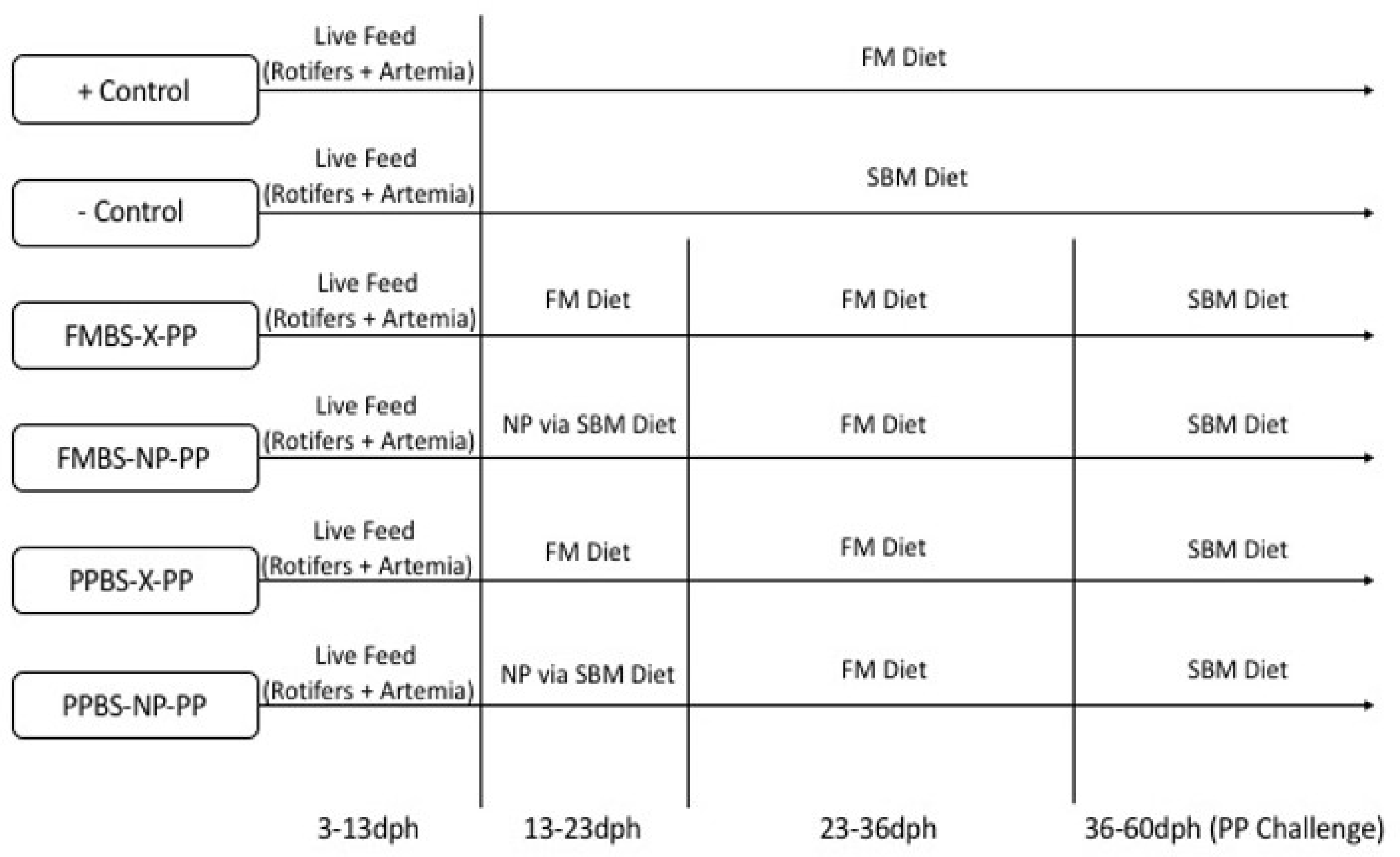
Feeding regime for each group.

### Sampling and Measuring

After the NP stage and the PP-Challenge stage of the study, five fish from each tank were sampled and stored in a 10% formalin solution for histology analyses. At the conclusion of the study, five fish in each tank were euthanized with MS-222 and intestines were harvested and stored in RNALater (Sigma-Aldrich, St. Louis, MO, USA) for future analysis of gene expression and an assessment of intestinal health.

Fish were weighed on a weekly basis beginning at 24 dph in order to measure biomass and adjust feeding rates. The average weight per fish of each group was measured at the end of the study, as was the percent weight gain of each group during PP challenge. Average weight was calculated for each tank by dividing the final biomass by the number of fish in the tank. The data shown for each group are average weight per fish for the 3 replicates of the group. Weight gain was calculated by subtracting the initial weight (start of the PP-Challenge) of the tank from the total weight at the conclusion of the study (PP- challenge). Weight gain was then divided by initial weight and multiplied by 100 to obtain weight gain percentage. Survival was assessed for the duration of the study (3-60 dph) by dividing the final number of fish in the tank by the initial number of fish (70) and multiplying that number by 100.

### Gene Expression Analysis

Intestinal samples were processed using TRIzol Reagent (Ambion, Foster City, CA, USA) and RNA was extracted and purified using the On-Column PureLink™ DNase Treatment (PureLink™ RNA Mini Kit and PureLink™ DNase, Invitrogen, Carlsbad, CA, USA) following the manufacturer’s instructions. Once purified, the nanograms/µl of each RNA sample was obtained using a spectrophotometer (Nanodrop 2000c, Thermo Fisher Scientific, Waltham, MA, USA). From this point, 2 µg of RNA from each sample was reverse transcribed using the High Capacity cDNA Reverse Transcription Kit (Applied Biosystems, Foster City, CA, USA) to obtain a 20 µl cDNA solution. The cDNA solutions were then added to a tube with 380 µl of water, to produce the cDNA sample for each tank. Gene expression of each cDNA sample was measured using a Bio-Rad (Hercules, CA, USA) CFX96™ Real-Time PCR System. Each qPCR reaction mixture (20 µl) contained 9 µl of cDNA sample, 10 µl of PowerUp™ SYBR™ Green Master Mix (Thermo Fisher Scientific, Waltham, MA, USA), and 0.5 µl of 800 nMol each of forward and reverse primers. Primers (Table 2) were synthesized by Integrated DNA Technologies (Coralville, IA, USA). Each qPCR reaction was run in technical duplicates. The qPCR cycle consisted of 95°C for 10 min, followed by 40 cycles of 95°C for 20 seconds and 60°C for 35 seconds, followed by a melting curve to ensure the amplification of only a single product in each well. Relative gene expression was calculated using the 2ΔΔCt method, normalizing the target gene expression to the expression of *ef1a* (reference gene) [5].

**Table 2.**
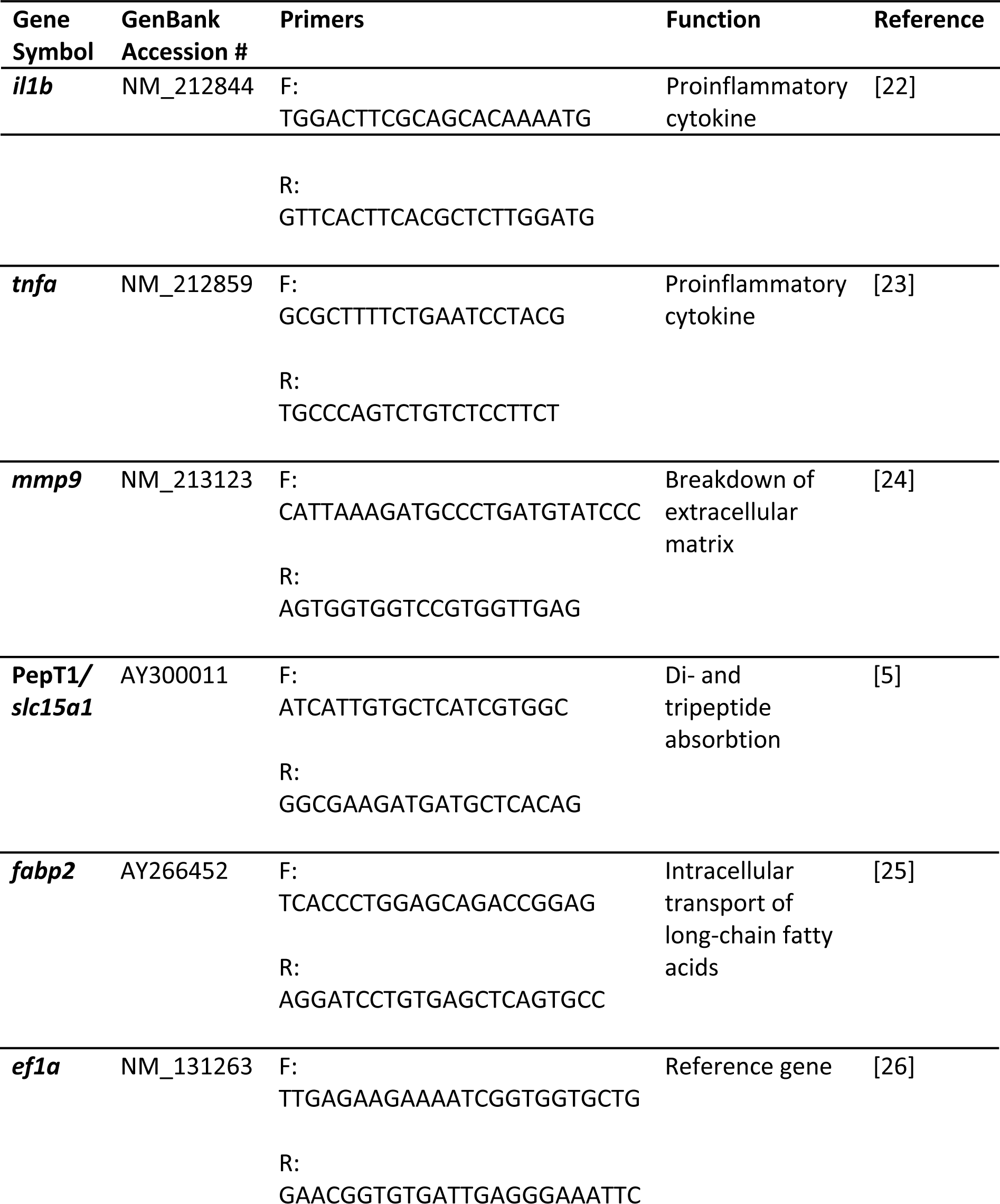
Primers used for gene expression analysis.

### Histological Analysis

Histological slides were prepared by Saffron Scientific Histology Services (Carbondale, IL). Intestines previously fixed in 10% neutral buffered formalin were processed to paraffin using a Sakura enclosed automated tissue processor (Netherlands). The three representative areas of zebrafish intestines were orientated for cross sections embedded together in the same block. Five-micrometer serial sections were cut with a Leica manual microtome (Buffalo Grover, IL) and placed on water bath at 44°C. Sections were placed on positive charged slides. After drying, slides were stained with hematoxylin and eosin and cover-slipped using acrylic mounting media. The histological analysis focused on the hind-gut portions of fish digestive tract. Pictures of each slide at 100X magnification were obtained using a microscope (Leica DMI 300B) and camera (Leica DMC 290) combination, with the software LAS V4.4 (Leica Camera, Wetzler, Germany). From these pictures, individual lengths and widths were taken of intact villi using ImageJ (NIH, Betheseda, MD, USA). Length and width data were measured in the distal portion of the intestine. Villus length was measured from the tip of the villus to the luminal surface, and villi width was measured across the base of the villus at the luminal surface. The length-to-width ratio of each villus was determined by dividing the length by the width.

### Statistical Analysis

Results are presented as means (± standard deviation). One-way ANOVA was used to test the data, and a Tukey test was run to test the differences between groups. Differences between groups are considered significant at *p* values < 0.05. Statistical analysis was run using R software.

## Results

### Growth performance

The (+) Control group had the highest average weight at the conclusion of the study, significantly higher than all other groups except FMBS-X-PP (Table 3). The PPBS-NP-PP group had the lowest average weight amongst the groups. There were no significant differences in weight gain during the PP-Challenge amongst the challenged groups, but both PPBS groups had a significantly lower weight gain than the (+) Control group. Survival did not significantly vary across the different treatment groups.

**Table 3.** Treatment effect on growth performance measures. Values are presented as means (± std. dev). Superscript letters indicate statistical significance between groups. The significance was determined using a One-Way ANOVA and a Tukey Test with a p value <0.05.

| Group | Avg. Weight (g) | Avg. Weight Gain during PP Challenge (%) | Survival (%) |
| --- | --- | --- | --- |
| + Control | 0.1913<sup>c</sup> (± 0.0109) | 271.71<sup>b</sup> (± 21.75) | 88.10 (± 0.82) |
| - Control | 0.1393<sup>ab</sup> (± 0.0009) | 232.50<sup>ab</sup> (± 3.85) | 83.81 (± 2.18) |
| FMBS-X-PP | 0.1676<sup>bc</sup> (± 0.0024) | 242.11<sup>ab</sup> (± 4.14) | 83.33 (± 6.44) |
| FMBS-NP-PP | 0.1467<sup>ab</sup> (± 0.0192) | 236.28<sup>ab</sup> (± 28.17) | 82.38 (± 8.73) |
| PPBS-X-PP | 0.1453<sup>ab</sup> (± 0.0114) | 221.10<sup>a</sup> (± 19.57) | 76.67 (± 7.33) |
| PPBS-NP-PP | 0.1315<sup>a</sup> (± 0.0093) | 228.80<sup>a</sup> (± 2.20) | 80.48 (± 14.31) |

### Histology

The first set of histological samples were taken prior to the start of the PP-Challenge, at 36 dph. In this set, distal intestine villus length-to width-ratio was significantly lower in all groups relative to the (+) Control ratio, except for the PPBS-X-PP group (Table 4). The second set of histological samples were taken after the PP-Challenge, at the conclusion of the study. In this set, the (+) Control and PPBS-NP-PP groups had the highest villus length to width ratios in the distal portion of the intestine, although it was only significantly higher than the villus length to width ratio of the FMBS-NP-PP group (Table 4).

**Table 4.**
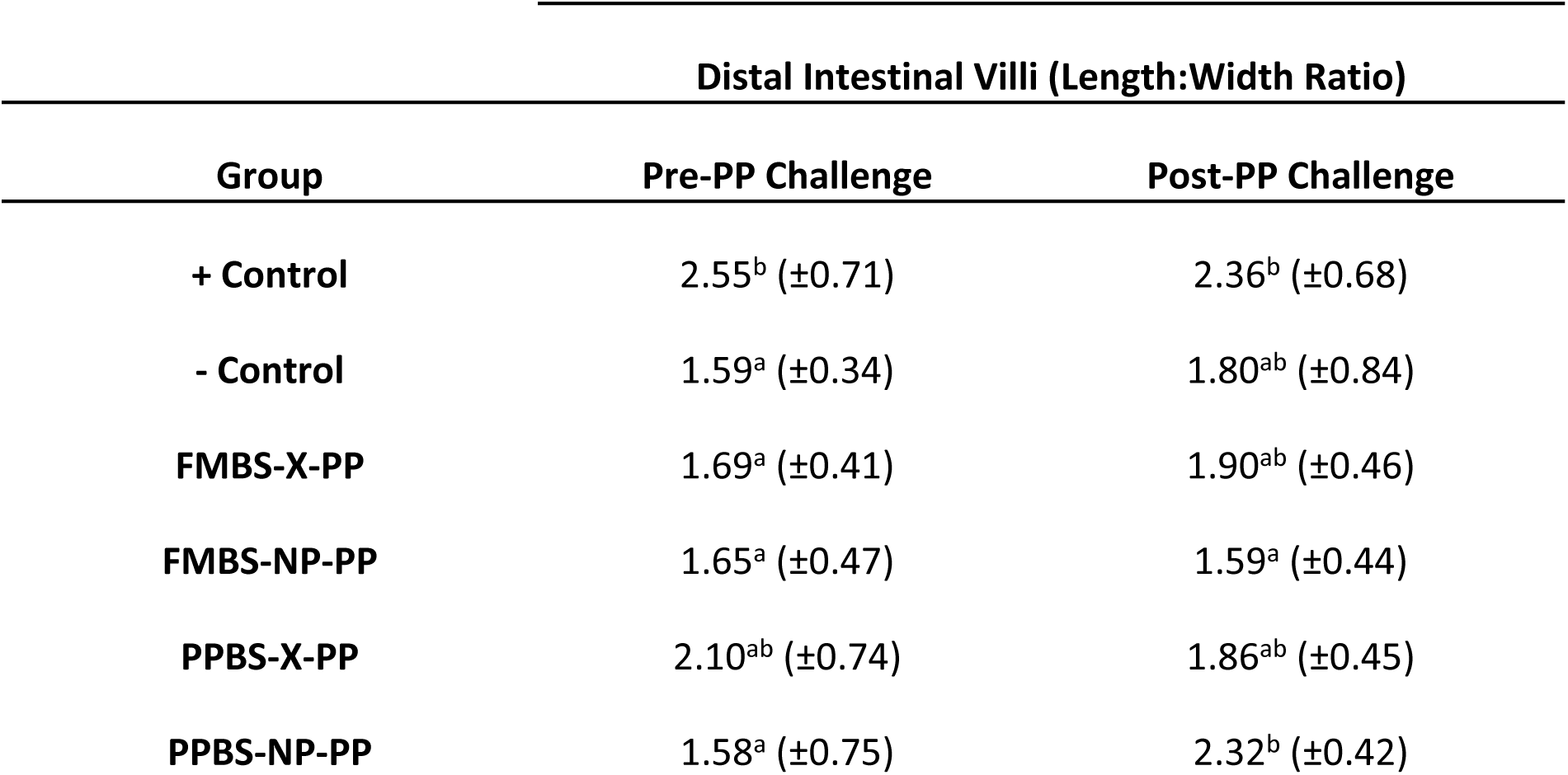
Treatment effect on the length-to-width ratio of villi in the distal portion of the intestine. Values are presented as means (± std. dev). Superscript letters indicate statistical significance between groups. The significance was determined using a One-Way ANOVA and a Tukey Test with a p value <0.05.

### Gene Expression

For four of the five genes investigated in this study, *il1b, tnfa, mmp9,* and *fabp2*, there were no significant differences in expression between groups (Fig 2). While not significant, the two NP groups seemed to have higher *fabp2* expression than the non-programmed groups from the same broodstock. However, treatment-specific differences were observed for expression of the Pept1 protein, responsible for the absorption of di- and tripeptides. Expression of PepT1 was lowest in the two groups that were offspring from the PP brood-stock, and expression in the PPBS-X-PP group was significantly lower expression than all other groups. Between the two PPBS groups, PepT1 expression was significantly higher in the group receiving nutritional programming (PPBS-NP-PP).

**Fig 2.**
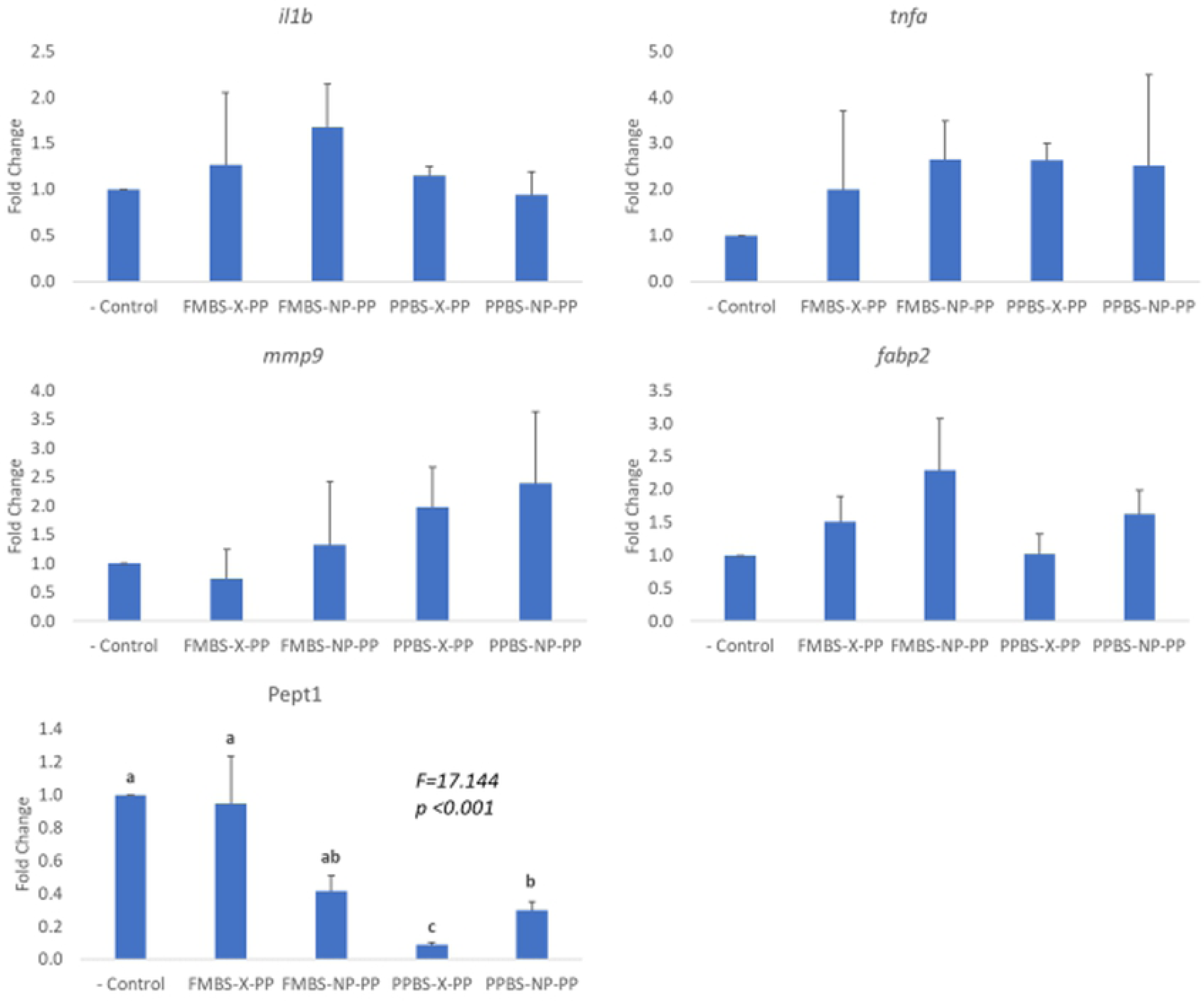
Intestinal gene expression. Relative gene expression of the group is represented as a fold change relative to the negative control group (fold change=1). Values provided are mean fold change + S.E.M (standard error of the mean). Results of One-way ANOVA and Tukey test are shown on graphs when significant. Significant differences (p<0.05) between groups are represented by different letters.

## Discussion

### Growth performance

While the significant programming effects reported in other studies [13, 17]were not observed in our growth performance study, it is important to note that weight gains among the FMBS were not significantly different. A similar programming study using yellow perch (*Perca flavescens*) also found no significance in growth rate between programmed and non-programmed fish [27]. In another study, NP with dietary carbohydrates from plant products initially improved growth performance in European sea bass (*Dicentrarchus labrax*) larvae; however, programming effects faded over time and no signifcant difference in growth was observed between programmed and nonprogrammed juveniles [28]. Interestingly, a similar study performed in our laboratory under the same conditions did find that NP improved growth performance of fish fed a SBM diet [29]. One possible explanation for the lack of programming influence in the present study could be species-specific differences. It has been found that different strains of fish can respond to plant protein differently [10,13,30,31], which may explain why significant programming effects on growth were not observed. The only significance detected in the weight gain data was that the two PPBS groups showed a decrease in weight gain compared to the (+) Control group which is in agreement with results obtained from other studies showing the negative effect of dietary PP on fish growth [1,2,4,7] The advantage of NP of broodstock lies in the fact that exposure to the potentially unfavorable nutritional trigger (in this case SBM) could induce an adaptation mechanism to the same trigger in the offspring without jeopardizing the growth of the broodstock or the quality of the gonads due to brief exposure time [17]. This is therefore applicable to fish species characterized by an extended gonad maturation period of several months. In zebrafish, gonad growth and maturation extends only to 2-4 weeks depending on the water conditions [32], and therefore the three-week exposure to SBM might have diminished the overall quality of both eggs and sperm and ultimately the fitness of the offspring even though supplemental feed in a form of *Artemia* nauplii was provided. The negative effect of PP feeding seemed to be reflected by low fecundity and hatching rates, which were not measured but observed by the researchers conducting the study. The decreased growth performance of the two PPBS groups observed in the study could also provide evidence that the parental dietary effect may have a larger impact on offspring performance than a NP effect of the broodstock or the offspring itself.

The lowest average weight per fish at the conclusion of the study was observed in the dual programmed group (PPBS-NP-PP), which was about 68% of the average weight in the (+) Control group. There could be a few factors affecting decreased weight in this group. In a related study done on Gilthead Seabream (*Sparus aurata*), researchers tested the effects of fish oil replacement with linseed oil in brood-stock diets on the performance of the progeny [17]. They found that while there were no signficant effects of partial fish oil replacement on growth of progeny, there was a significant decrease in the final body weight of offspring from broodstock that received the diet without any fish oil [17]. Similarly in our study, full replacement of fishmeal in the broodstock diet could have limited the growth potential of fish in the next generation. Some studies have indicated that poor growth response can be improved when fish are fed a plant-based diet at an early age regardless of their genetic capacity for growth [13]. However, our results are more in an agreement with a different study; these researchers argued that “fish that grow faster on fishmeal diet are likely to grow faster on plant protein-based diets” [33]. We postulate that fish that have a “better start” because of enhanced broodstock nutrition (fishmeal vs. SBM) might perform better regardless of the diet received at a later stage. In fact, the negative impact that broodstock nutrition might have on fish reproduction has been well reported [18,27,34–37]. For instance, soybean meal inclusion in goldfish (*Carrasius auratus*) diets disrupted sex hormone biosynthesis and caused a reduction in fertilization and hatching rates [37]. Other studies showed that feeding broodstock a high replacement level of fishmeal and fish oil significantly reduced larval growth and also juvenile growth, so it is apparent that the effects can persist for an extended period of time [18]. Thus, it is feasible that while both broodstocks had the same nutritional history leading up to the study, the three weeks of SBM-diet prior to breeding may have had enough of an impact on the reproductive processes as well as gamete quality in the PPBS to affect the offspring. This is supported by a recent study finding that, regardless of nutritional history (SBM vs fishmeal), reproductive quality of broodstock is not affected as long as they are fed a fishmeal-based diet during gametogenesis [27]. This finding consequently suggests that NP of broodstock should be adjusted to the specific reproductive characteristics of fish species, including the time required for gonad maturation.

### Intestinal health

The intestinal villus length-to-width ratio provides an important measure of the efficiency of nutrient absorption within the intestine. A higher villus length-to-width ratio indicates increased surface area for nutrient absorption, which correlates with an improved specific growth rate [10]. Reduced villus length to width ratio in the distal intestine has previously been found to be a result of intestinal inflammation caused by SBM, and is associated with a decrease in the absorption capacity of enterocytes in the intestine [1,10,12]. Our study found significantly improved villus length-to-width ratio of the PPBS-NP-PP group. Prior to the PP-Challenge, the PPBS-NP-PP group had the lowest villus length to width ratio (1.58 ±0.75). At the conclusion of the study the villus length to width ratio was 2.32 (±0.42), higher than all other groups that experienced the PP-Challenge and significantly higher than the FMBS-NP-PP group. The reasoning behind this improvement may lie in the adaptability of the hindgut to prolonged exposure to SBM. In a similar study done on common carp (*Cyprinus carpio*), an omnivorous member of the Cyprinidae, after four to five weeks of exposure to dietary SBM the distal portion of the intestine started to recover morphologically from SBM-induced inflammation [12]. The PPBS-NP-PP group had dietary SBM as an environmental stimulus through the broodstock programming and through traditional NP induced during early development of the offspring. Our results suggest that the dual programming of the PPBS-NP-PP group could potentially improve the villus length-to-width ratio of the distal intestine and improve nutrient absorption in the intestine when fed a SBM-based diet.

The *il1b*, *tnfa*, and *mmp9* genes were measured to observe the level of inflammation within the intestine. The levels of *il1b* and *tnfa* are directly related to intestinal inflammation because they are proinflammatory cytokines and their increased expression marks an immune response within the intestine [5]. The presence of the proinflammatory cytokines can also trigger the upregulation of *mmp9,* a gene involved in the breakdown of the extracellular matrix [5,38,39]. An increase in *mmp9* signals an increase in the breakdown of tissue within the intestine resulting from inflammation [5,38,39]. We hypothesized that through NP induced in broodstock, or early development of offspring, or both, we could reduce the level of intestinal inflammation caused by SBM and we would observe a down-regulation of each of these genes compared to the (–) Control group. However, our results showed that for each of the three genes, expression levels did not significantly vary among groups, contradicting results obtained by Perera and Yufera [5]. A key reason for this discrepancy could be the length of the actual programming and the induction timing. Perera and Yufera [5] had a NP period of 3 days starting right after the mouth opening compared to the 10 days of programming after the live food feeding in our study. Another reason for the varying results could be the diet formulations. Perera and Yufera [5], tested 100% SBM and 100% soy protein concentrate (SPC) diets separately. They found that the SPC diet had no significant effect on the inflammatory cytokines in the intestine due to lack of anti-nutritional factors, while the SBM diet did. A key factor between the two diets is soy saponin, which was found to be the driver of intestinal inflammation in fish fed a SBM-based diet [1]. Soy saponin is found in SBM but not in SPC, so the 15% inclusion of SPC in the diet for this study may have reduced the saponin levels in the diet and resulted in lower levels of inflammatory cytokines.

While there were few trends amongst the inflammatory genes, the genes associated with nutrient absorption did reveal some significant results. The two genes measured to assess the nutrient absorption levels within the intestines were PepT1 and *fabp2* [5]*. Fabp2* is responsible for encoding an intracellular fatty acid-binding protein that is involved in the transport of long-chain fatty acids [40]. While no significant differences between the groups were observed, there was a key trend. In both the FMBS and PPBS groups, the group that received NP through exposure to SBM in early age showed numerically higher level of *fabp2*. Perera and Yufera [5], observed a similar trend where programming with SBM led to increased expression of fatty acid-binding proteins providing support for the idea behind using NP as a means for possible improvement of dietary lipid absorption.

The most significant results in this study were seen in the expression of PepT1. Increased expression of peptide transporters in the intestine is a key factor in the absorption of protein components in fish [41, 42]. Our results showed that the two PPBS groups had much lower expression of PepT1 than the FMBS groups, with the PPBS-X-PP group having a significantly lower expression than any other group (only ∼ 9% compared to the (–) Control group). A previous study [43], found that, during a starved state, intestinal PepT1 expression is decreased in fish. It is possible that NP of the broodstock with SBM reduced PepT1 expression and that effect was further passed down to the offspring through potential epigenetic change in that PepT1 gene. Lower expression of peptide transporter could indicate that somehow SBM-based diet was not fully utilized and perhaps led to fish reaching a physiological state of starvation. This result could possibly suggest that short exposure to dietary SBM during gonad development and maturation in zebrafish might not necessarily be positive for the broodstock stage itself, in contrast with results from other studies [17, 18]. Furthermore, PepT1 expression in PPBS-NP-PP fish was significantly higher compared to the PPBS-X-PP group. The significant impact of NP on PepT1 possibly suggests that the mechanism of NP in fish is related to alteration of genes involved in nutrient uptake and transportation. Previous studies have shown that increased expression of genes involved in nutrient utilization can be significantly correlated with an increase in growth performance [9, 10]. Venold et al. [10], found that higher expression of *fabp2* correlated with a higher specific growth rate in rainbow trout. While the impact on growth performance in this study was not observed, it is important to note the duration of the feeding experiment. This study lasted only 60 dph to ensure that the sexual dimorphism of mature zebrafish would not impact the growth results. It is plausible to assume, however, that if the study lasted longer the increased *fabp2 or* PepT1 expression may have lead to an increased growth performance as seen in other fish species. Further studies should therefore focus on the association between NP, expression of genes involved in nutrient absorption, and growth in other fish species.

## Conclusion

Although the hypothesized programming effects did not produce a significant increase in the growth performance of fish fed SBM diet neither through the broodstock nor through early exposure of the offspring (or both), findings from this study can be key to the future improvement of SBM utilization in aquaculture feeds. The histological and gene expression results support the use of programming as a means of improving nutrient absorption through increased intestinal villus surface area and the upregulation of nutrient (peptide) absorption genes. Future studies are encouraged on other fish species to further explore the impact of NP on the epigenome of the broodstock and its offspring to identify which other genes are modulated to better understand the mechanism of NP and its potential to improve utilization of feeds based on lower-quality raw materials.

## Acknowledgements

We thank Saffron Scientific Histology Services (Carbondale, IL) for providing the histological services for this study.

## Author Contributions

Experimental conception and design: KK, MW. Experiment management and execution: GSM, KK, MW. Data analyses: GSM, VJM, GT, SR. Writing (original draft): GSM. Writing (review and edit): GSM, KK, VJM, GT.

